# Clusters of sub-Saharan African countries based on sociobehavioural characteristics and associated HIV incidence

**DOI:** 10.1101/620450

**Authors:** Aziza Merzouki, Janne Estill, Erol Orel, Kali Tal, Olivia Keiser

**Author notes:** <u>Corresponding author:</u> Aziza Merzouki, Institute of Global Health, University of Geneva, Chemin des Mines 9, 1202 Geneva, Switzerland.

## Abstract

**Introduction:** HIV incidence varies widely between sub-Saharan African (SSA) countries. This variation coincides with a substantial sociobehavioural heterogeneity, which complicates the design of effective interventions. In this study, we investigated how sociobehavioural heterogeneity in sub-Saharan Africa could account for the variance of HIV incidence between countries.

**Methods:** We analysed aggregated data, at the national-level, from the most recent Demographic and Health Surveys of 29 SSA countries [2010-2017], which included 594’644 persons (183’310 men and 411’334 women). We preselected 48 demographic, socio-economic, behavioural and HIV-related attributes to describe each country. We used Principal Component Analysis to visualize sociobehavioural similarity between countries, and to identify the variables that accounted for most sociobehavioural variance in SSA. We used hierarchical clustering to identify groups of countries with similar sociobehavioural profiles, and we compared the distribution of HIV incidence (estimates from UNAIDS) and sociobehavioural variables within each cluster.

**Results:** The most important characteristics, which explained 69% of sociobehavioural variance across SSA among the variables we assessed were: religion; male circumcision; number of sexual partners; literacy; uptake of HIV testing; women’s empowerment; accepting attitude toward people living with HIV/AIDS; rurality; ART coverage; and, knowledge about AIDS. Our model revealed three groups of countries, each with characteristic sociobehavioural profiles. HIV incidence was mostly similar within each cluster and different between clusters (median(IQR); 0.5/1000(0.6/1000), 1.8/1000(1.3/1000) and 5.0/1000(4.2/1000)).

## Introduction

The burden of HIV in sub-Saharan Africa (SSA) is the heaviest in the world; in 2017, 70% of HIV-infected people lived in this region (“Fact sheet - World AIDS Day 2018”). However, HIV prevalence and incidence vary widely between SSA countries. The region is heterogeneous and sociobehavioural and cultural factors vary widely within and between countries, complicating the design of effective interventions. This heterogeneity ensures that no “one-size-fits-all” approach will stop the epidemic. This is why WHO (“Global Health Sector Strategy on HIV 2016-2021: Towards Ending AIDS”) highlights the need to use data and numerical methods to tailor interventions to specific populations and countries based on quantitative evidence.

So far, studies of HIV risk factors or risk factors for the uptake of interventions against HIV have generally been limited to specific sub-populations (Sangowawa & Owoaje, 2012; Kidman & Anglewicz, 2016; Ashaba et al., 2018), sub-national regions (Bailey et al., 2007; Gray et al., 2007; Eaton et al., 2014; Pons-Duran et al., 2016) or single countries (Antelman et al., 2007; Gregson et al., 2010; Tsai & Venkataramani, 2015; Lakew, Benedict & Haile, 2015; Kelly, Weiser & Tsai, 2016; Smith Fawzi et al., 2016; Kim et al., 2016; McGillen et al., 2018; Merzouki et al., 2020). Recent studies included up to 31 SSA countries, but narrowly focused their inquiries to examine, for example, the association between socioeconomic inequalities (Hajizadeh et al., 2014), high-risk sexual behaviour (Kenyon, Buyze & Schwartz, 2018), or HIV-related stigma (Chan & Tsai, 2015; Kelly, Weiser & Tsai, 2016) with HIV testing, treatment uptake, ART (antiretroviral treatment) adherence, or HIV prevalence. Most used standard statistical methods like descriptive statistics (Sangowawa & Owoaje, 2012; Smith Fawzi et al., 2016), linear or logistic regression (Mondal & Shitan, 2013; Delavande, Sampaio & Sood, 2014; Chan & Tsai, 2015; Kidman & Anglewicz, 2016; Ashaba et al., 2018), or concentration indices (Hajizadeh et al., 2014; Pons-Duran et al., 2016; Kim et al., 2016), to assess health inequity and the impact of a small number of variables (5 to 13) on the HIV epidemic. In a previous work, we also investigated the associations and possible causal relationships between sociobehavioural factors at the individual-level that are potentially related to the risk of acquiring HIV in 29 SSA countries using Bayesian Network models (Baranczuk et al., 2019). But, these methods do not inform us on how HIV risk factors vary across SSA and which characteristic sociobehavioural patterns at the country-level are actually associated with different rates of new HIV infections in the region.

Recent studies have shown that unsupervised learning and clustering analysis allows us to find hidden sub-groups of people with varying drivers and potentially different risk levels of having or acquiring HIV(Engl, Smittenaar & Sgaier, 2019; Merzouki et al., 2020). At the country-level, comparing and characterising SSA countries would allow us to test the hypothesis that sociobehavioural heterogeneity might account for spatial variance of HIV epidemic, and inform effective country-specific interventions.

In this study, we used dimensionality reduction and unsupervised machine learning techniques (Principal Component Analysis and hierarchical clustering) to identify the most important factors of 48 national attributes that might account for the variability of HIV incidence across SSA, and identified the sociobehavioural profiles that characterized different levels of HIV incidence, based on Demographic and Health Survey (“The DHS Program - Quality information to plan, monitor and improve population, health, and nutrition programs”) data from 29 SSA countries that were aggregated at the national-level.

## Methods

### Data

We used aggregated data from the most recent Demographic and Health Surveys (DHS) of 29 SSA countries completed between 2010 and 2017, that were available up to July 2018 (**Table S1**). DHS typically gathers nationally representative(“DHS Sampling and Household Listing Manual (English)”) data on health (including HIV-related data) and population (including social, behavioural, geographic and economic data) every 5 years. These data are publicly accessible at the individual-, district- and country-level. We used data aggregated at the national-level.

We attempted to include in our analysis all variables possibly influencing the risks of HIV acquisition among population aged 15-49 years that were available for the 29 SSA countries through the StatCompiler tool (“STATcompiler”). We only included variables that varied significantly across the region, and did not strongly correlate with other variables. We finally pre-selected the following variables: age (under 25 vs older); rurality (rural vs urban); religion (Christian, Muslim, Folk/Popular religions, unaffiliated, others); marital status (married or in union vs widowed/divorced/other), number of wives for men (1, ≥2) or co-wives for women (0, 1, ≥2); literacy (literate vs illiterate); media access (with access to newspaper, television and radio at least once a week vs without such access); employment (worked in the last 12 months and currently working vs others); wealth (Gini coefficient); age at first sexual intercourse (first sexual intercourse by age 15 vs older); general fertility (number of births to women of reproductive age in the last 3 years); contraception use (using any method of contraception vs not using any); condom use (belief that a woman is justified in asking condom use if she knows her husband has a Sexually Transmitted Infection (STI) vs belief that she is not justified); number of sexual partners in lifetime; unprotected higher risk sex (men who had sex with a non-marital, non-cohabiting partner in the last 12 months and did not use condom during last sexual intercourse vs not); paid sex (men who ever paid for sexual intercourse vs never paid for sex); unprotected paid sex (men who used condom during the last paid sexual intercourse in the last 12 months vs did not use condom); gender-based violence (wife beating justified for at least one specific reason vs not justified for any reason); married women participation to decision making (yes vs no); gender of household head (female vs male); comprehensive correct knowledge about AIDS (yes vs no); HIV testing (ever receiving an HIV test vs never tested); male circumcision (yes vs no); ART coverage (i.e. percentage of people on antiretroviral treatment among those living with HIV); and accepting attitudes toward people living with HIV/AIDS (would buy fresh vegetables from a shopkeeper with AIDS vs would not); see **Table 1** for a complete summary of the variables.

**Table 1 -.**
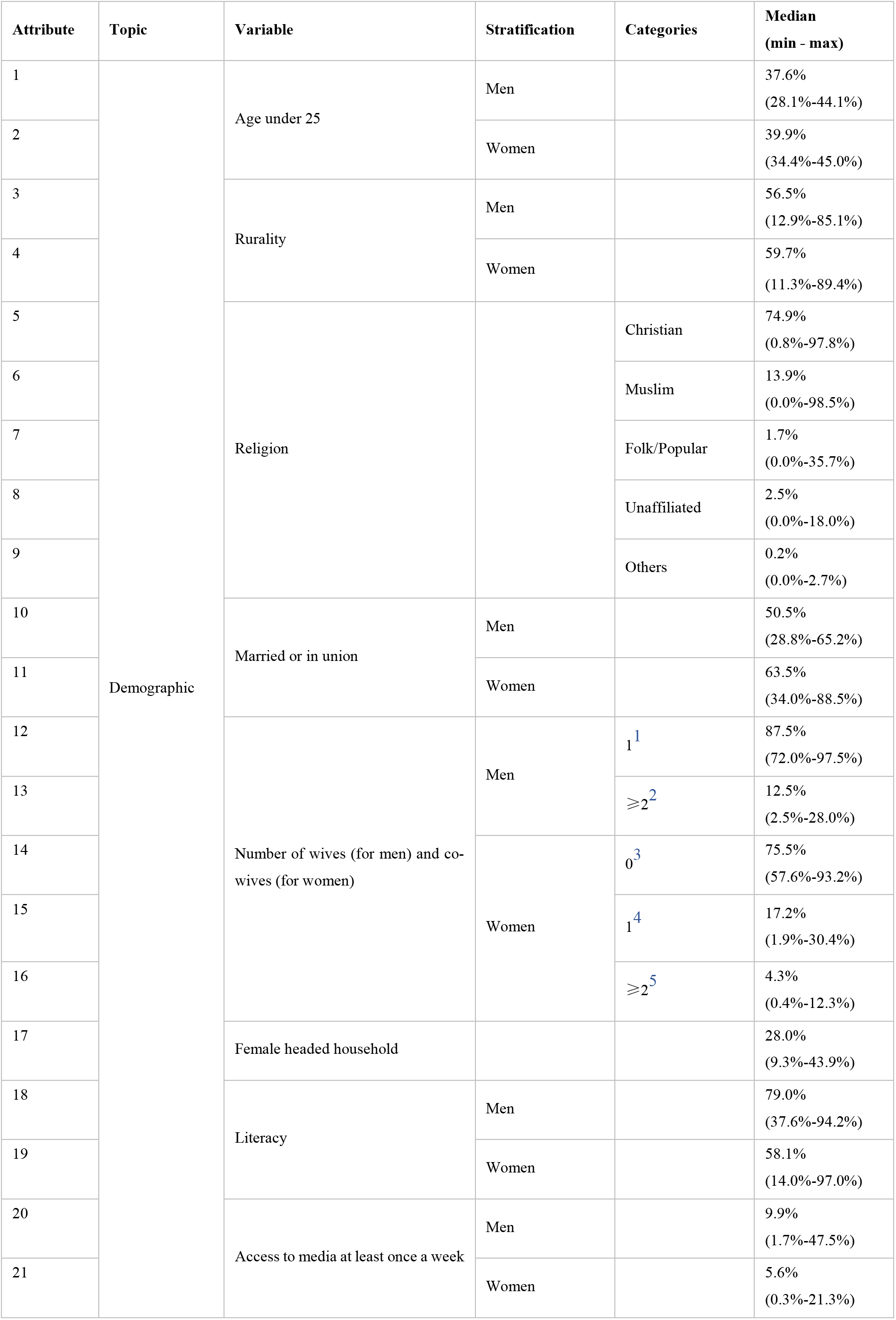
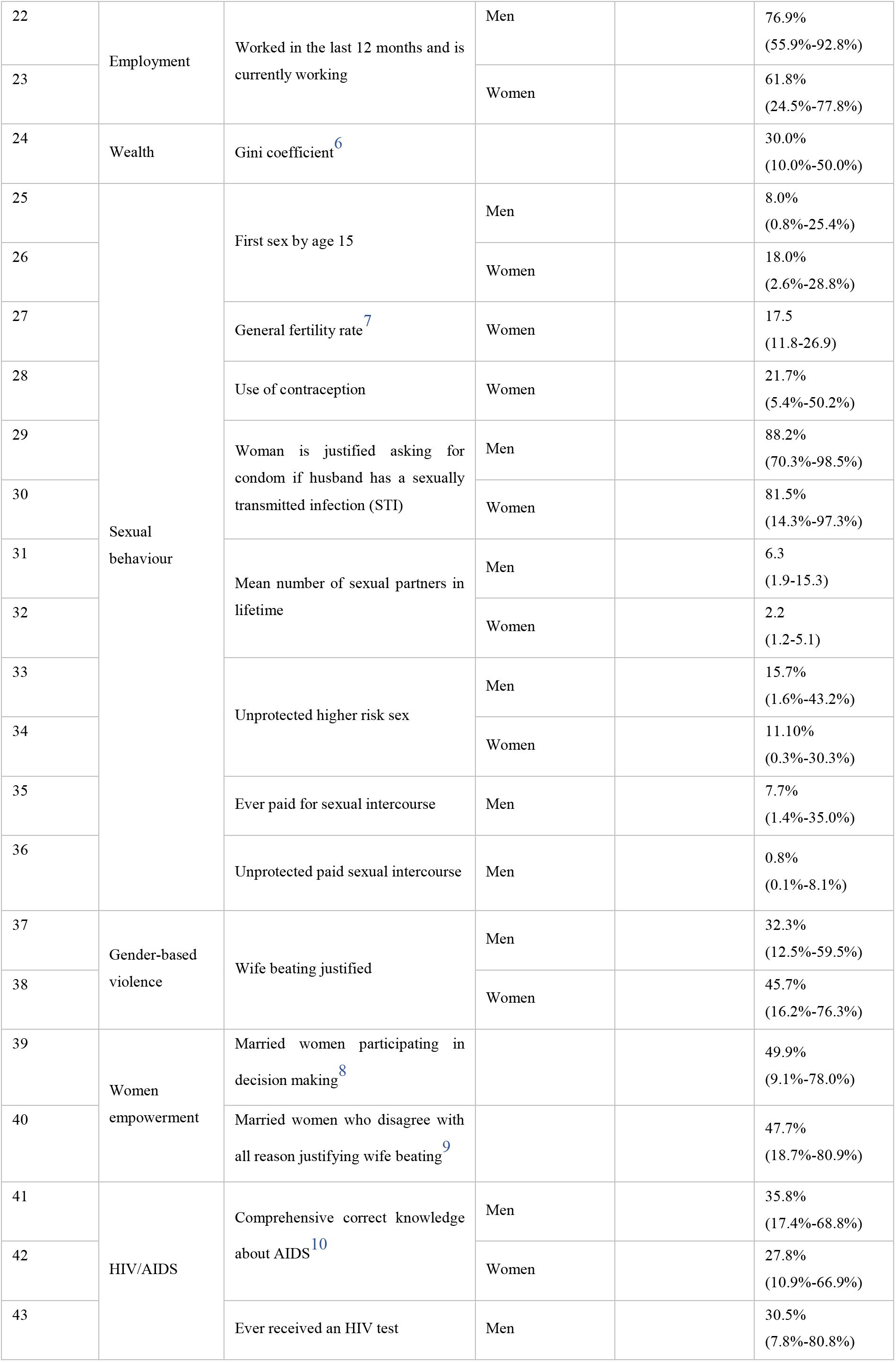
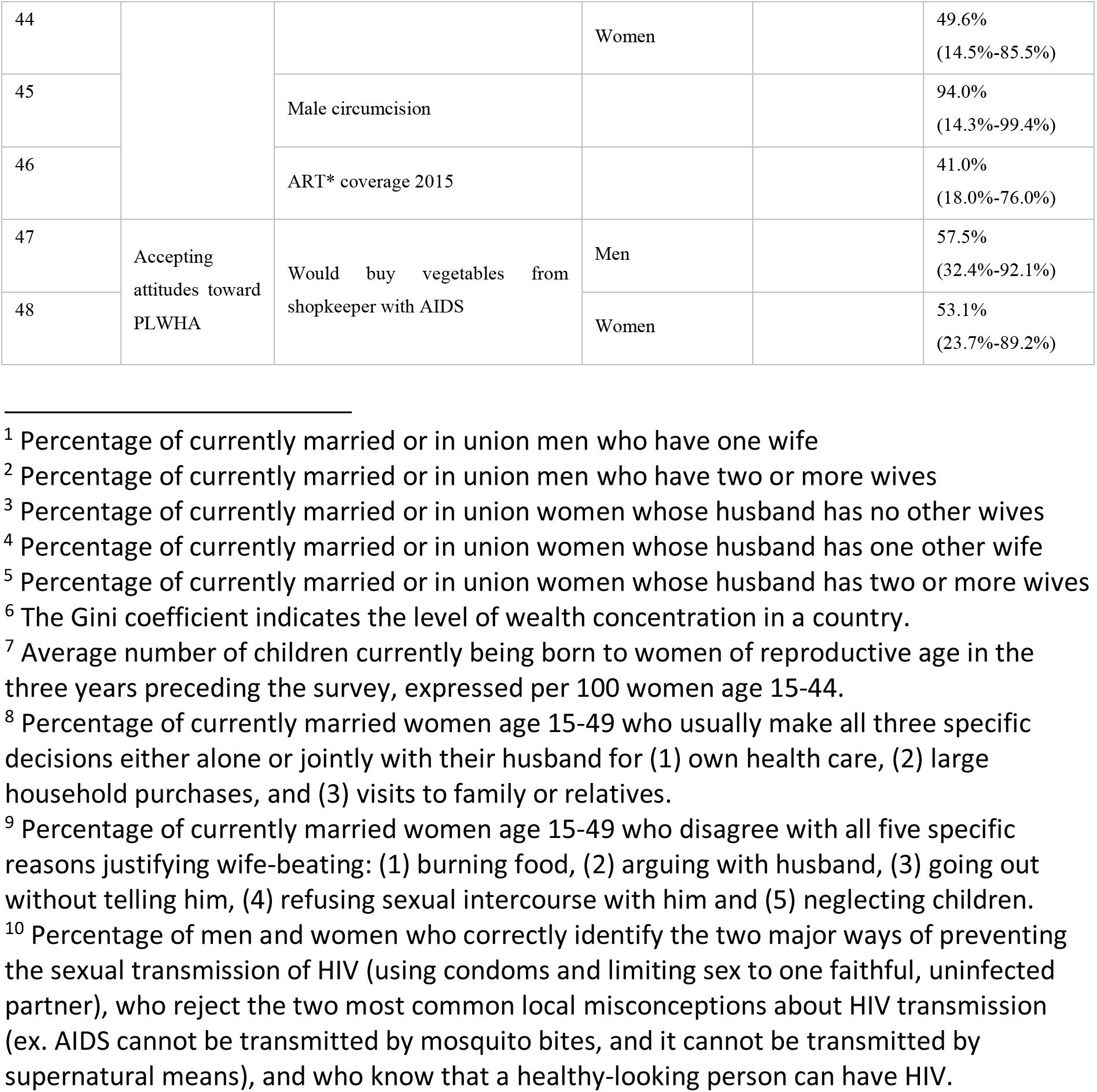
Socio-economic and behavioural variables included in the analysis. Median values across all 29 countries are shown with the minimum and maximum values. *ART = antiretroviral therapy.

| Attribute | Topic | Variable | Stratification | Categories | Median<br>(min - max) |
| --- | --- | --- | --- | --- | --- |
| 1 | Demographic | Age under 25 | Men |  | 37.6%<br>(28.1%-44.1%) |
| 2 |  |  | Women |  | 39.9%<br>(34.4%-45.0%) |
| 3 |  | Rurality | Men |  | 56.5%<br>(12.9%-85.1%) |
| 4 |  |  | Women |  | 59.7%<br>(11.3%-89.4%) |
| 5 |  | Religion |  | Christian | 74.9%<br>(0.8%-97.8%) |
| 6 |  |  |  | Muslim | 13.9%<br>(0.0%-98.5%) |
| 7 |  |  |  | Folk/Popular | 1.7%<br>(0.0%-35.7%) |
| 8 |  |  |  | Unaffiliated | 2.5%<br>(0.0%-18.0%) |
| 9 |  |  |  | Others | 0.2%<br>(0.0%-2.7%) |
| 10 |  | Married or in union | Men |  | 50.5%<br>(28.8%-65.2%) |
| 11 |  |  | Women |  | 63.5%<br>(34.0%-88.5%) |
| 12 |  | Number of wives (for men) and co-wives (for women) | Men | 1<sup>1</sup> | 87.5%<br>(72.0%-97.5%) |
| 13 |  |  |  | ≥2<sup>2</sup> | 12.5%<br>(2.5%-28.0%) |
| 14 |  |  | Women | 0<sup>3</sup> | 75.5%<br>(57.6%-93.2%) |
| 15 |  |  |  | 1<sup>4</sup> | 17.2%<br>(1.9%-30.4%) |
| 16 |  |  |  | ≥2<sup>5</sup> | 4.3%<br>(0.4%-12.3%) |
| 17 |  | Female headed household |  |  | 28.0%<br>(9.3%-43.9%) |
| 18 |  | Literacy | Men |  | 79.0%<br>(37.6%-94.2%) |
| 19 |  |  | Women |  | 58.1%<br>(14.0%-97.0%) |
| 20 |  | Access to media at least once a week | Men |  | 9.9%<br>(1.7%-47.5%) |
| 21 |  |  | Women |  | 5.6%<br>(0.3%-21.3%) |
| 22 | Employment | Worked in the last 12 months and is currently working | Men |  | 76.9%<br>(55.9%-92.8%) |
| 23 |  |  | Women |  | 61.8%<br>(24.5%-77.8%) |
| 24 | Wealth | Gini coefficient<sup>6</sup> |  |  | 30.0%<br>(10.0%-50.0%) |
| 25 | Sexual behaviour | First sex by age 15 | Men |  | 8.0%<br>(0.8%-25.4%) |
| 26 |  |  | Women |  | 18.0%<br>(2.6%-28.8%) |
| 27 |  | General fertility rate<sup>7</sup> | Women |  | 17.5<br>(11.8-26.9) |
| 28 |  | Use of contraception | Women |  | 21.7%<br>(5.4%-50.2%) |
| 29 |  | Woman is justified asking for condom if husband has a sexually transmitted infection (STI) | Men |  | 88.2%<br>(70.3%-98.5%) |
| 30 |  |  | Women |  | 81.5%<br>(14.3%-97.3%) |
| 31 |  | Mean number of sexual partners in lifetime | Men |  | 6.3<br>(1.9-15.3) |
| 32 |  |  | Women |  | 2.2<br>(1.2-5.1) |
| 33 |  | Unprotected higher risk sex | Men |  | 15.7%<br>(1.6%-43.2%) |
| 34 |  |  | Women |  | 11.10%<br>(0.3%-30.3%) |
| 35 |  | Ever paid for sexual intercourse | Men |  | 7.7%<br>(1.4%-35.0%) |
| 36 |  | Unprotected paid sexual intercourse | Men |  | 0.8%<br>(0.1%-8.1%) |
| 37 | Gender-based violence | Wife beating justified | Men |  | 32.3%<br>(12.5%-59.5%) |
| 38 |  |  | Women |  | 45.7%<br>(16.2%-76.3%) |
| 39 | Women empowerment | Married women participating in decision making<sup>8</sup> |  |  | 49.9%<br>(9.1%-78.0%) |
| 40 |  | Married women who disagree with all reason justifying wife beating<sup>9</sup> |  |  | 47.7%<br>(18.7%-80.9%) |
| 41 | HIV/AIDS | Comprehensive correct knowledge about AIDS<sup>10</sup> | Men |  | 35.8%<br>(17.4%-68.8%) |
| 42 |  |  | Women |  | 27.8%<br>(10.9%-66.9%) |
| 43 |  | Ever received an HIV test | Men |  | 30.5%<br>(7.8%-80.8%) |
| 44 |  |  | Women |  | 49.6%<br>(14.5%-85.5%) |
| 45 |  | Male circumcision |  |  | 94.0%<br>(14.3%-99.4%) |
| 46 |  | ART* coverage 2015 |  |  | 41.0%<br>(18.0%-76.0%) |
| 47 | Accepting<br>attitudes toward<br>PLWHA | Would buy vegetables from<br>shopkeeper with AIDS | Men |  | 57.5%<br>(32.4%-92.1%) |
| 48 |  |  | Women |  | 53.1%<br>(23.7%-89.2%) |

<sup>6</sup> The Gini coefficient indicates the level of wealth concentration in a country.
<sup>7</sup> Average number of children currently being born to women of reproductive age in the three years preceding the survey, expressed per 100 women age 15-44.
<sup>8</sup> Percentage of currently married women age 15-49 who usually make all three specific decisions either alone or jointly with their husband for (1) own health care, (2) large household purchases, and (3) visits to family or relatives.
<sup>9</sup> Percentage of currently married women age 15-49 who disagree with all five specific reasons justifying wife-beating: (1) burning food, (2) arguing with husband, (3) going out without telling him, (4) refusing sexual intercourse with him and (5) neglecting children.
<sup>10</sup> Percentage of men and women who correctly identify the two major ways of preventing the sexual transmission of HIV (using condoms and limiting sex to one faithful, uninfected partner), who reject the two most common local misconceptions about HIV transmission (ex. AIDS cannot be transmitted by mosquito bites, and it cannot be transmitted by supernatural means), and who know that a healthy-looking person can have HIV.

We represented each country using 48 dimensions. Each dimension corresponded to an attribute in **Table 1**, such as the percentage of women married or in union, the mean number of sexual partners in a lifetime for men, the percentage of Christian populations and the Gini coefficient in this country. Data were represented as percentages; the mean number of sexual partners in lifetime was scaled using min-max normalisation. Most of these country-level data were exported from the DHS with the StatCompiler tool, except for data on religion that we obtained from Pew-Templeton Global Religious Futures Project (“Religions in Africa | African Religions | PEW-GRF”), and ART coverage that we obtained from UNAIDS’ AIDSinfo (“AIDSinfo | UNAIDS”). We used the latest (2018) UNAIDS estimates of national HIV incidence for the year 2016 (“AIDSinfo | UNAIDS”; “Estimates Methods 2018”).

### Ethics approval

The study was conducted using aggregated data that are publicly accessible through StatCompiler (for DHS data), Pew-Templeton Global Religious Futures Project, and UNAIDS’ AIDSinfo. No ethics approval was needed from our side.

### Analysis

We used Principal Component Analysis (PCA) (Trevor Hastie, Robert Tibshirani & Jerome Friedman; James et al., 2013) to reduce the dimensionality of data from 48 to two dimensions (2D) so we could visualise sociobehavioural similarity between SSA countries. PCA allows to extract directions, called principal components (PCs), along which the variation of data is maximal. The first two PCs, which explain the most variance, represent the axes of the 2D-space used for visualisation. Countries closest to each other when projected on the 2D-space correspond to similar countries in terms of demographic, socioeconomic and behavioural characteristics. PCs consist of a linear combination of the initial 48 dimensions and can therefore be interpreted in terms of the original sociobehavioural variables (“PCA - Principal Component Analysis Essentials - Articles - STHDA”). We can then identify the variables among those we included in the analysis, that explain most of the sociobehavioural heterogeneity across SSA.

We identified groups of similar SSA countries in terms of sociobehavioural characteristics using hierarchical clustering. Pairwise country dissimilarity was calculated using the Euclidian distance (**Equation S1**). These distances were used by the hierarchical clustering algorithm to create a *dendrogram* with 29 terminal nodes representing the countries to be grouped. Cutting the dendrogram at a certain height produces clusters of similar countries. The number of clusters depends on the height at which the tree is cut. We measured the quality of the clustering results using the Silhouette Index, and selected the optimal number of clusters such that it maximised this index (**Equation S4**).

We compared the HIV incidence of countries between the identified clusters, and visualised the distribution of HIV incidence within each cluster using *box plots*. We also identified the sociobehavioural variables that characterized the resulting clusters, and visualized the distribution of these variables within each cluster using *density plots*.

We used the open source R language, version 3.5.1 for our analysis. Code and country-level data are available on GitLab (https://gitlab.com/AzizaM/dhs_ssa_countries_clustering), in two separate folders “code” and “data_countries”, respectively.

## Results

We analysed aggregated data from surveys that included 594,644 persons in total, 183,310 men and 411,334 women(“DHS Sampling and Household Listing Manual (English)”), ranging from 9,552 in Lesotho to 56,307 in Nigeria. Adult HIV incidence ranged from 0.14/1000 in Niger to 19.7/1000 in Lesotho in 2016. HIV prevalence ranged from 0.4% in Niger to 23.9% in Lesotho (**Table S1**). Sociobehavioural characteristics varied widely between SSA countries (**Table 1**).

### Visualizing the SSA countries: Geographical and sociobehavioural similarities

Using PCA, we found that the first principal component (PC) explained 49.5% and the second 19.5% of the total sociobehavioural variance across SSA among the 48 variables we considered (**Figure 1**). The original sociobehavioural variables that contributed most to these PCs were religion (12.6% for Muslim and 12.1% for Christian populations), male circumcision (9.4%), number of sexual partners (7.8% for men and 3.4% for women), literacy (6.1% for women and 3.2% for men), HIV testing (5.5% for men and 5.4% for women), women’s participation in decision making (3.8%), an accepting attitude towards those living with HIV/AIDS (3.6% for women and 3.2% for men), rurality (3.0% for women and 2.7% for men), ART coverage (2.5%), and women’s knowledge about AIDS (2.5%) (**Figure 1B** and **Figure S1**).

**Figure 1 -.**
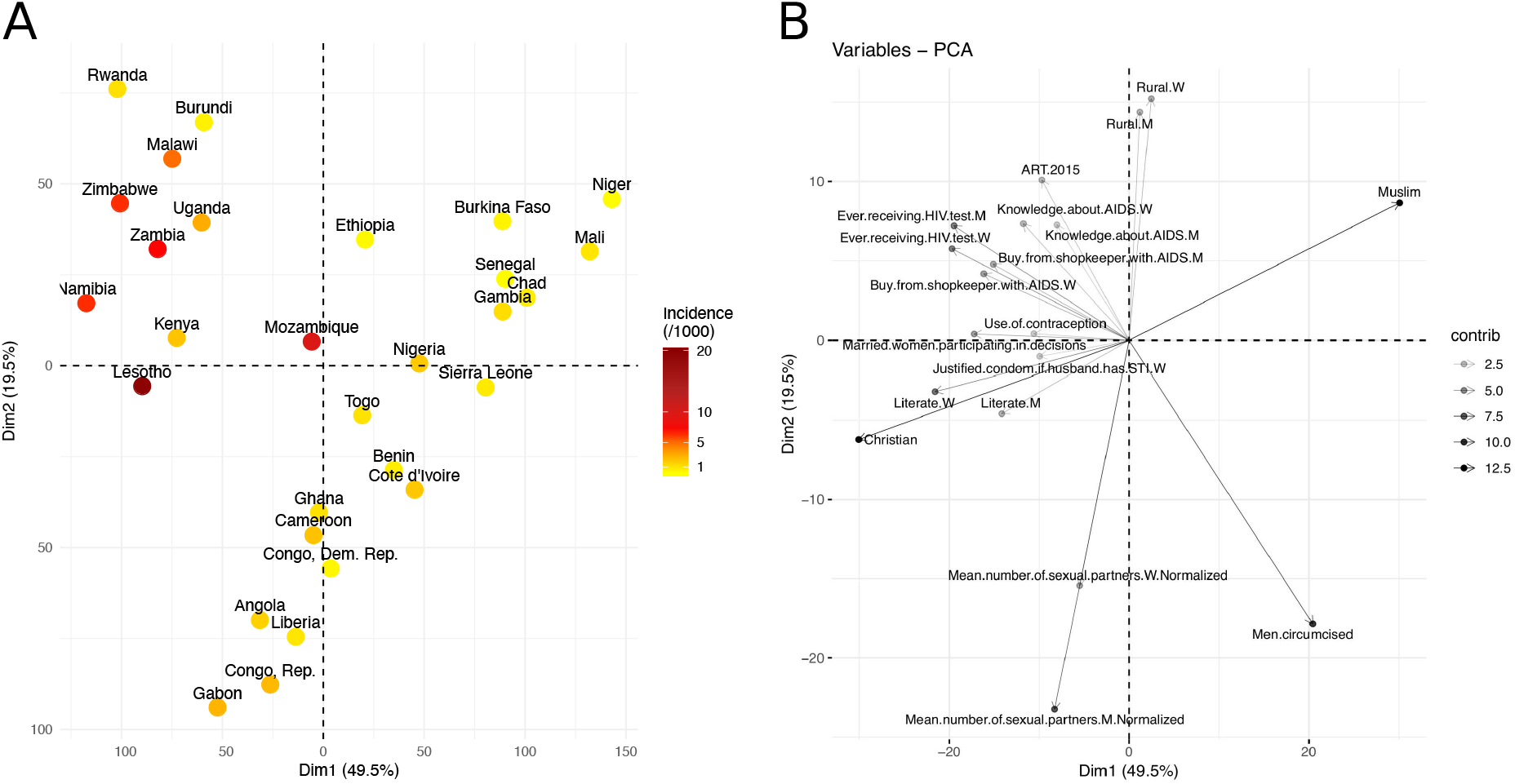
Visualization of the sociobehavioural similarity between SSA countries using PCA. **(A) Projection of the SSA countries on a 2D-space, based on their socio-economic and behavioural factors.** The two dimensions (first two PCs), Dim1 and Dim2, explained 69% of the variance in the data. Countries are coloured based on their HIV incidence per 1000 population (15-49) in 2016. **(B) Correlation plot of the original variables with the first and second dimensions (Dim1, Dim2).** The variable transparency represents its contribution (in %) to the two dimensions. Moving along a variable’s vector leads toward a region of the 2D-space where the variable levels tend to be higher, e.g. upper right quadrant contains mainly Muslim countries, while upper left quadrant contains countries with higher levels of HIV testing and knowledge about AIDS.

Projecting the 29 SSA countries in two dimensions produced a roughly V-shaped scatterplot (**Figure 1A**). **Figure 1B** shows how the 48 original sociobehavioural variables vary over the 2D-space. At the end of the V-shape’s left branch, Eastern and Southern African countries, such as Namibia, Zimbabwe, Malawi, Zambia and Uganda, lied next to each other. In these countries, fewer men are circumcised, but the percentage of literate people who had accepting attitudes toward people living with HIV/AIDS (PLWHA) was higher and so was the uptake of HIV testing. Knowledge about AIDS and ART coverage were also high. The end of the right branch, in the upper right quadrant, included countries from the Sahel region, like Senegal, Burkina Faso, Mali, Niger and Chad, where the percentage of Muslims is higher and people have fewer sexual partners. The lower tip of the V-shape included countries from West and Central Africa, like Liberia, Ghana, Côte d’Ivoire, Democratic Republic of the Congo, and Gabon, where people have more sexual partners, more men are circumcised, and the population is less rural.

### Clustering the SSA countries and analysis of the associated HIV incidence

The hierarchical clustering of the 29 SSA countries produced a dendrogram (**Figure 2A)**. Cluster compactness and separation were optimal (maximum silhouette index = 0.3) when we cut the dendrogram at a height that separated countries into three groups (**Figure 2B**).

**Figure 2 -.**
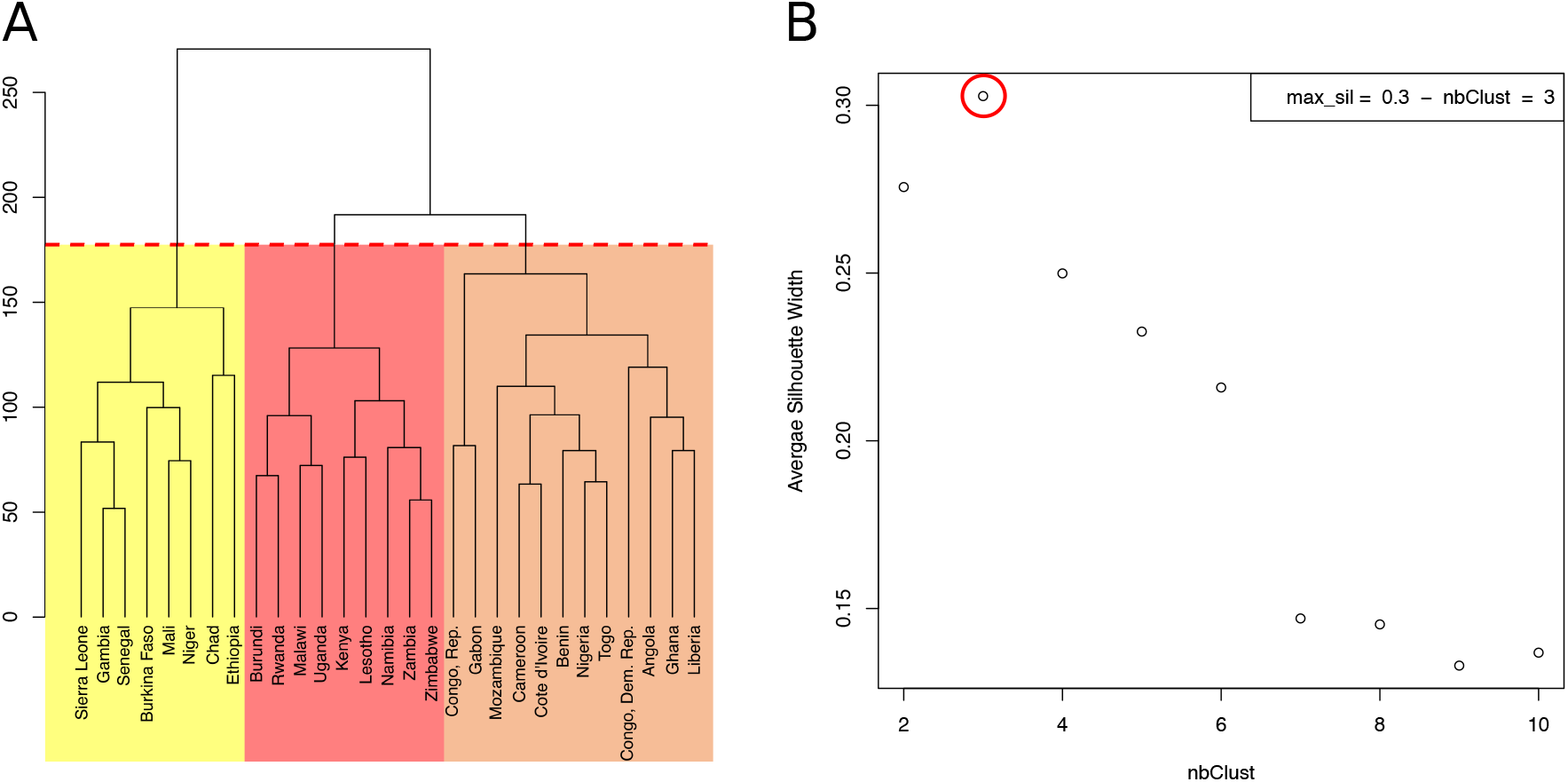
Hierarchical clustering of 29 sub-Saharan African countries. **(A) Dendrogram.** Cutting the tree at the height of the red dashed line results in three clusters, highlighted in yellow, orange and red. **(B) Average Silhouette width for different numbers of clusters**. The number of clusters (X axis), from 2 to 10, corresponds to different heights at which the dendrogram was cut. The maximum average Silhouette width was obtained for 3 clusters (red circle).

The countries of the first cluster, in yellow, had the lowest HIV incidence (median of 0.5/1000 population) (**Figure 3**). This cluster included countries from the Sahel Region, where the population was mostly rural (median of 71.1% for men) and Muslim (median of 86.2%). On the one hand, countries of this cluster were characterized by many factors that could account for low HIV incidence and prevalence. Countries were characterized by high proportions of circumcised men (median of 95.0%), high percentages of women who were married or lived in union (median of 70.6%), late sexual initiation for men (median of 1.9% of men who had their first sexual intercourse by the age of 15), low numbers of sexual partners (median of 3.5 partners for men), low percentages of unprotected higher-risk sex (median of 9.7% for men) and low percentages of men having ever paid for sex (median of 3.9%). Polygyny (Reniers & Tfaily, 2012; Eaton et al., 2014), an institutionalized form of sexual concurrency, was also frequent in this region (median of 22.3%). On the other hand, this cluster was also characterized by frequent belief that wife beating is justified (median of 61.2% for women), and low levels of literacy (median of 29.0% for women). Participation of married women in decision making (median of 18.5%), contraceptive prevalence (median of 13.9%), and knowledge about AIDS (median of 23.7% for women) was also low. These countries had low percentages of people ever tested for HIV (median of 19.2% for men; 36.6% for women), low ART coverage (median of 38.0%) and low levels of acceptance of PLWHA (Median of 47.4% for men); see **Figure 4** and **Figure 5**.

**Figure 3 -.**
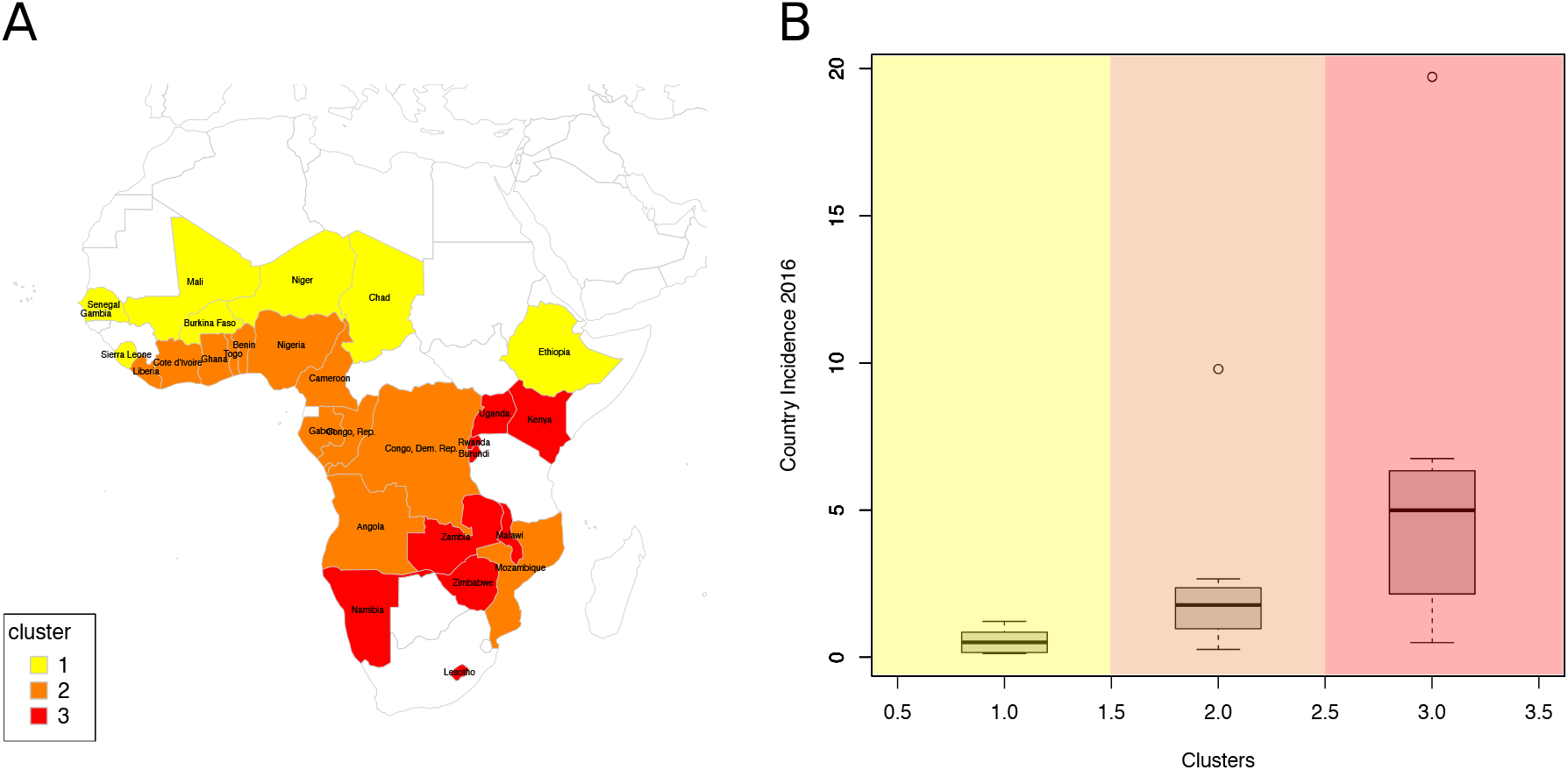
Analysis of the resulting clusters. **(A) Map of clustered sub-Saharan countries.** Countries are coloured based on the cluster to which they belong. **(B) Box plots of the HIV incidence distribution within each cluster**.

**Figure 4 -.**
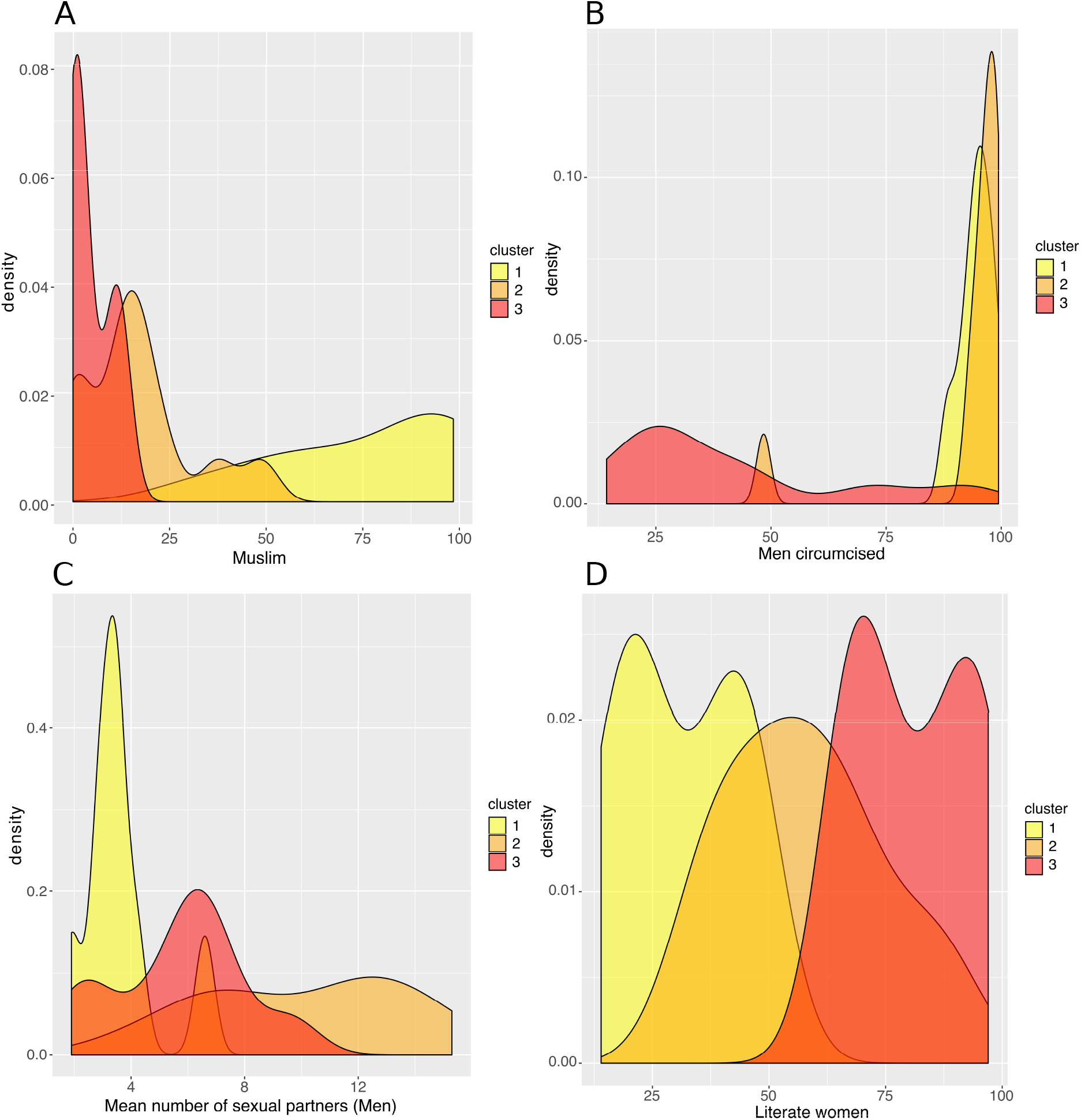
Analysis of the resulting clusters in terms of their sociobehavioural characteristics. Density plots per cluster of (A) the percentage of Muslim population, (B) the percentage of circumcised men, (C) the mean number of sexual partners in a man’s lifetime, (D) the percentage of literate women, (E) the percentage of men who have ever received an HIV test, (F) the percentage of men who say they would buy fresh vegetables from a vendor whom they knew was HIV+, (G) the percentage of women with a comprehensive knowledge about AIDS and (H) the ART coverage in 2015.

**Figure 5 -.**
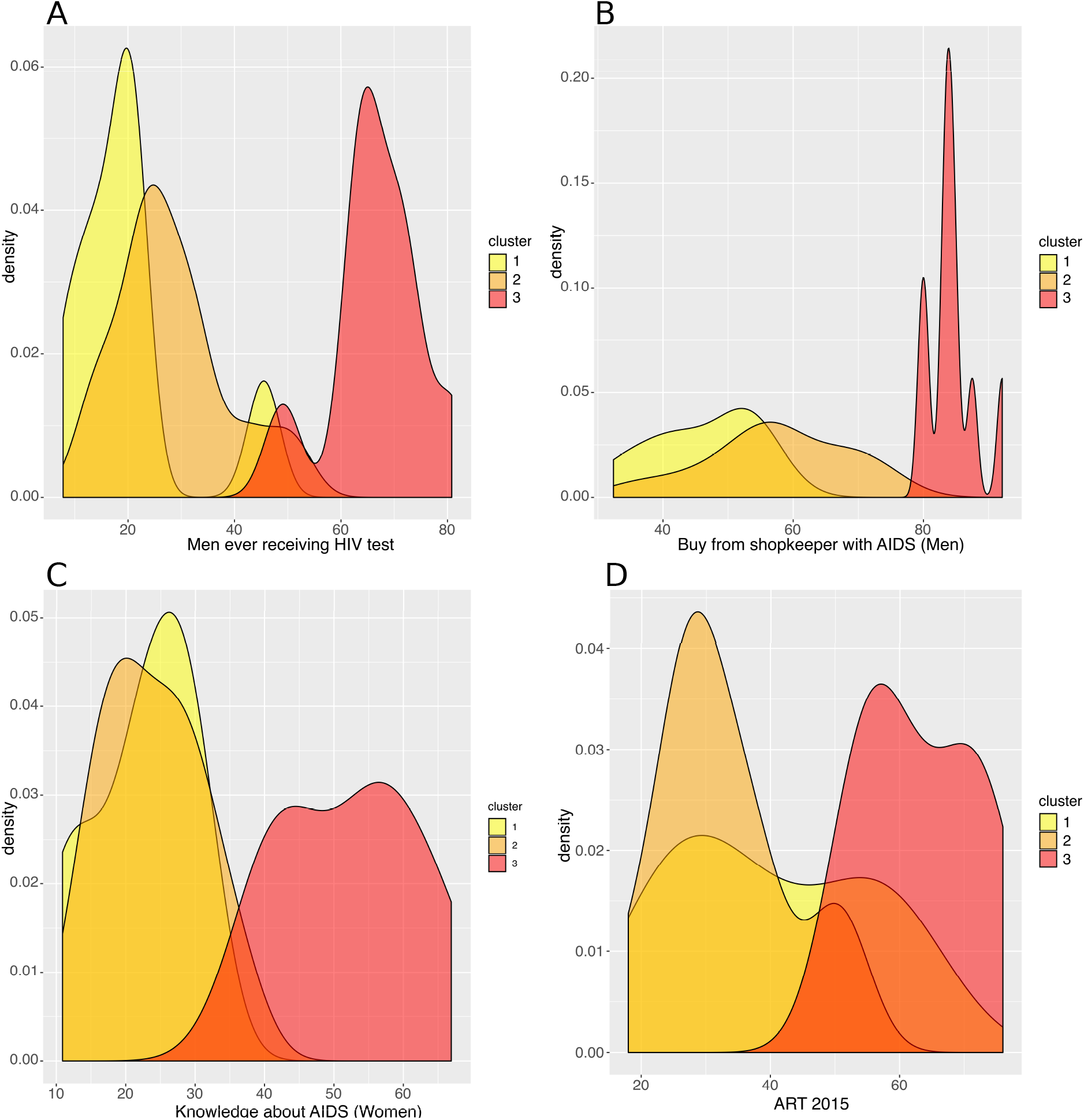
Analysis of the resulting clusters in terms of their HIV-related attributes. Density plots per cluster of (A) the percentage of men who have ever received an HIV test, (B) the percentage of men who say they would buy fresh vegetables from a vendor whom they knew was HIV+, (C) the percentage of women with a comprehensive knowledge about AIDS and (D) the ART coverage in 2015.

The countries of the second cluster, coloured in orange, included countries from West and Central Africa. These countries had a rather low HIV incidence (median of 1.8/1000 population), though Mozambique was a remarkable outlier, with a high HIV incidence (9.8/1000 population) (**Figure 3**). Like the first cluster, these countries had a high percentage of circumcised men (median of 97.0%, except in Mozambique where only 48.4% of men were circumcised). However, these countries were also characterized by the lowest proportions of rural populations (median of 49.0% for men), the highest numbers of sexual partners (median of 10.1 for men), early sexual initiation (median of 12.0% of men who had their first sexual intercourse by the age of 15), and more frequent unprotected high-risk sex (median of 24.3% for men) and paid sexual intercourse (median of 9.5% for men). HIV testing uptake (median of 25.8% for men and 48.6% for women), knowledge about AIDS (median of 23.6% for women), and ART coverage (median of 31.0%) were all low.

The third cluster, in red, included Southern and East African countries. These countries had high HIV incidence (median of 5.0/1000 population), except two countries that had a lower HIV incidence: Rwanda (1.1/1000 population) and Burundi (0.5/1000) (**Figure 3**). Countries belonging to the third cluster were characterized by the lowest percentage of circumcised men (median of 27.9%). But they were also the ones with the highest uptake of HIV testing (median of 65.2% for men; 83.3% for women) and ART (median of 61.0%), and the highest percentage with knowledge about AIDS (median of 54.6% for women) and accepting attitudes towards PLWHA (median of 84.4% for men). This cluster was also characterized by the highest percentage of literacy (median of 80.2% for women), high use of contraceptives (median of 42.6%), low percentages of unprotected high-risk sex (median of 9.8% for men) and higher percentages of married women participating in decision making (median of 67.7%) and women-headed households (median of 31.0%). Rwanda and Burundi had the lowest HIV incidence and were characterized by a lower number of sexual partners (Rwanda, 2.6; Burundi, 2.1) vs a median of 6.3 partners for men in the other countries of the third cluster. They also had larger per capita rural populations (Rwanda, 80.4%; Burundi, 89.4%) vs a median of 61.3% for women in the other countries of the same cluster.

## Discussion

Using hierarchical clustering, we identified the most important characteristics that explained 69% of the sociobehavioural variance among the variables we assessed in SSA. The variables that contributed the most to the sociobehavioural heterogeneity across the 29 SSA countries and among the 48 attributes we included in this analysis are religion, number of sexual partners, literacy, HIV testing, women’s participation in decision making, accepting attitude towards those living with HIV/AIDS, rurality, ART coverage, and women’s knowledge about AIDS. We found three groups of countries with similar sociobehavioural patterns, and HIV incidence was also similar within each cluster.

In the first cluster, PLWHA were not widely accepted, and the population had an overall low-level knowledge about AIDS. Stigma may be more widespread in this region and explain the lower uptake of interventions among people who are HIV-positive. The relatively low number of people who are living with HIV lowers the general public’s exposure to this group and may increase stigma (Chan & Tsai, 2017). Stigma can also result from cultural and religious beliefs that link HIV/AIDS with sexual transgressions, immorality and sin (Campbell et al., 2005; Mbonu, van den Borne & De Vries, 2009).

We think that the apparent contradiction between the presence of many high-risk factors and the low HIV incidence in most countries of the second cluster could be explained by the high proportion of circumcised men. In line with this theory, Mozambique, the only country in this cluster with very high HIV prevalence and incidence, had few circumcised men. Previous observational studies and trials have confirmed the protective effect of male circumcision (Bailey et al., 2007; Gray et al., 2007; Lei et al., 2015; Sharma et al., 2018).

Countries of the third cluster, with the highest HIV incidence, were also the ones with the highest knowledge about AIDS (Chan & Tsai, 2017), ART coverage, uptake of HIV testing, and with the most accepting attitudes toward PLWHA. They also had the lowest percentage of unprotected higher risk sex. These findings are consistent with earlier studies that found broad ART coverage may reduce social distancing towards PLWHA and HIV-related stigma in the general population(Chan & Tsai, 2015; Chan, Tsai & Siedner, 2015). Reduced social distancing and stigma is associated with higher uptake of voluntary HIV counselling and testing (Kalichman, 2003; Kelly, Weiser & Tsai, 2016), and less sexual risk-taking among HIV positive people (Delavande, Sampaio & Sood, 2014).

The high HIV incidence in Mozambique could be caused by any combination of the following factors: a high number of sexual partners; a low level of male circumcision; and a low level of literacy and knowledge about AIDS. The latter two factors can also be responsible for low uptake of HIV testing and ART. However, many West and Central African countries with population characteristics like Mozambique in terms of sexual practices, literacy, knowledge about AIDS, HIV testing and ART coverage, had much lower HIV prevalence and incidence, possibly because males were circumcised at twice the rate. Compared to other West and Central African countries, an aggravating factor in Mozambique can also be the high prevalence of female genital schistosomiasis(Hotez et al., 2019; Yegorov et al., 2019). On the other side of the spectrum, despite a low uptake of male circumcision, it is also possible that the combination of lower numbers of sexual partners, higher per capita rural populations, more literacy, more accurate knowledge about AIDS, more HIV testing, and broader ART coverage could account for lower HIV incidence, like in Rwanda and Burundi.

In contrast to more classical regression techniques that usually estimate the relation between independent sociobehavioural factors and a single outcome, dimensionality reduction and clustering techniques allow us to visualize the sociobehavioural heterogeneity across SSA countries, and identify characteristic patterns of factors that are shared by groups of countries and are found to be associated with different levels of HIV incidence. These results provide policy makers with a quick and clearer overview of sociobehavioural factors across SSA, and help identify targeted interventions that can reduce HIV transmission in a specific country or group of countries.

The cross-sectional nature of our data makes it impossible to determine precedence and causality between the sociobehavioural characteristics we measured and HIV prevalence and incidence. But the associations we identified can open lines of inquiry for researchers. Our study had the advantage of allowing us to compare countries and regions (clusters of countries), but ecological studies that use aggregated data are prone to confounding and ecological fallacy (Levin). Africa is an exceedingly diverse continent with many distinct sub-populations, so a study based on national population averages cannot explain HIV variation within countries. Recent studies have shown the complex geographical variation of the HIV epidemic and its drivers within Eastern and Southern African countries (Cuadros et al., 2017; Bulstra et al., 2020). Therefore, we intend to repeat our study at a lower level of granularity, using regional-, district- and individual-level data to capture differences within countries and learn more about the complex patterns of sociobehavioural factors that affect the sub-populations that are most at risk. We believe that machine learning is a promising innovative way to further explore and improve our understanding of these questions, in parallel to more classical techniques.

Our work has some other limitations. We used model estimates for HIV incidence, which may diverge from reality (Nsanzimana et al., 2017). Sociobehavioural data from the DHS are self-reported, therefore potentially biased; for instance, literate individuals may be more reluctant to admit accepting gender-based violence or having higher risk sex. And even though we included many more variables than is common practice (Hajizadeh et al., 2014; Lakew, Benedict & Haile, 2015; Kidman & Anglewicz, 2016; Kim et al., 2016; Ashaba et al., 2018; Kenyon, Buyze & Schwartz, 2018), we still had to exclude many factors that can play a critical role in HIV transmission, including the prevalence of excessive alcohol consumption, STIs (Looker et al., 2017; Torrone et al., 2018) (ex. genital herpes, gonorrhea and syphilis), endemic infections (Hotez et al., 2019; Yegorov et al., 2019)(ex. malaria and helminthiases), ART adherence and drug resistance, as well as the distribution of different HIV strains and subtypes (Buonaguro, Tornesello & Buonaguro, 2007; Esbjörnsson et al., 2019; Gartner et al., 2020). Some of the latter variables were not available from the DHS or were missing for at least one country included in our analysis.

## Conclusions

Our use of PCA allowed us to identify the most important characteristics among the variables we assessed that explained 69% of the sociobehavioural variance in SSA countries. With hierarchical clustering, we captured complex patterns of sociobehavioural characteristics shared by countries with similar HIV incidence, suggesting that the combination of sociobehavioural factors plays a key role in determining the course of the HIV epidemic. Our findings can help to design and predict the effect of targeted country-specific interventions to impede HIV transmission.

## Authors’ contributions

AM, JE and OK designed the study. AM and OK reviewed and selected the variables to be included in the analysis. AM wrote the code and performed the analysis with support from JE. EO reviewed analysis and code, and checked results reproducibility. AM interpreted the results with support from OK and KT. AM, OK and KT reviewed the literature. AM wrote the manuscript, which was reviewed by OK, JE, and KT. All authors have read and approved the final manuscript.

## Acknowledgements

We thank Zofia Baranczuk for helpful discussions.

