## Supplementary Material for "Clusters of sub-Saharan African countries based on sociobehavioural characteristics and associated HIV incidence"

#### Countries, DHS data and HIV incidence and prevalence

**Table S1- List of 29 sub-Saharan countries with their estimated HIV incidence (age 15-49) and prevalence (age 15-49) in 2010 and 2016** (UNAIDS's 2018 estimates *[1]* produced by Spectrum software).

| ID | Country | DHS year | HIV incidence per 1000 population (15-49) | | HIV prevalence (15-49) (%) | |
|---|---|---|---|---|---|---|
| | | | 2010 | 2016 | 2010 | 2016 |
| 1 | Angola | 2015-16 | 2.13 | 1.68 | 1.8 | 1.9 |
| 2 | Benin | 2011-12 | 0.69 | 0.63 | 1.1 | 1.0 |
| 3 | Burkina Faso | 2010 | 0.37 | 0.39 | 1.0 | 0.8 |
| 4 | Burundi | 2016-17 | 0.46 | 0.51 | 1.5 | 1.1 |
| 5 | Cameroon | 2011 | 2.98 | 2.21 | 4.6 | 3.8 |
| 6 | Chad | 2014-15 | 0.85 | 0.65 | 1.6 | 1.3 |
| 7 | Congo, Rep. | 2011-12 | 2.70 | 2.53 | 3.2 | 3.2 |
| 8 | Congo, Dem. Rep. | 2013-14 | 0.41 | 0.28 | 1.2 | 0.7 |
| 9 | Cote d'Ivoire | 2011-12 | 2.03 | 2.03 | 3.6 | 2.8 |
| 10 | Ethiopia | 2016 | 0.19 | 0.21 | 1.4 | 1 |
| 121 | Gabon | 2012 | 3.30 | 2.67 | 4.2 | 4.3 |
| 12 | Gambia | 2013 | 1.79 | 1.23 | 2.1 | 1.7 |
| 13 | Ghana | 2014 | 1.27 | 1.07 | 2.0 | 1.7 |
| 14 | Kenya | 2014 | 3.46 | 2.16 | 5.6 | 5.0 |
| 15 | Lesotho | 2014 | 22.00 | 19.70 | 22.2 | 23.9 |
| 16 | Liberia | 2013 | 1.05 | 0.92 | 2.0 | 1.5 |
| 17 | Malawi | 2015-16 | 8.70 | 5.00 | 10.9 | 9.9 |
| 18 | Mali | 2012-13 | 1.07 | 0.95 | 1.4 | 1.3 |
| 19 | Mozambique | 2011 | 14.53 | 9.79 | 13.3 | 12.7 |
| 20 | Namibia | 2013 | 9.06 | 6.34 | 13.0 | 12.3 |
| 21 | Niger | 2012 | 0.20 | 0.14 | 0.5 | 0.3 |
| 22 | Nigeria | 2013 | 2.32 | 1.89 | 3.2 | 2.8 |
| 23 | Rwanda | 2014-15 | 1.47 | 1.12 | 3.2 | 2.8 |
| 24 | Senegal | 2017 | 0.22 | 0.14 | 0.6 | 0.4 |
| 25 | Sierra Leone | 2013 | 1.16 | 0.78 | 1.6 | 1.4 |
| 26 | Togo | 2013-14 | 6.56 | 2.97 | 3.0 | 2.2 |
| 27 | Uganda | 2016 | 9.78 | 6.75 | 6.7 | 6.1 |
| 28 | Zambia | 2013-14 | 11.00 | 6.33 | 12.3 | 11.8 |
| 29 | Zimbabwe | 2015 | 2.13 | 1.68 | 15.0 | 13.7 |

### Variable contribution to Principle Components PC1 and PC2

**Figure S1- Variables sorted by their contribution to the first two Principle Components***[2]***, which explain 69% of the sociobehavioural variance between SSA countries**. Only variables that contributed more than 1% are displayed.

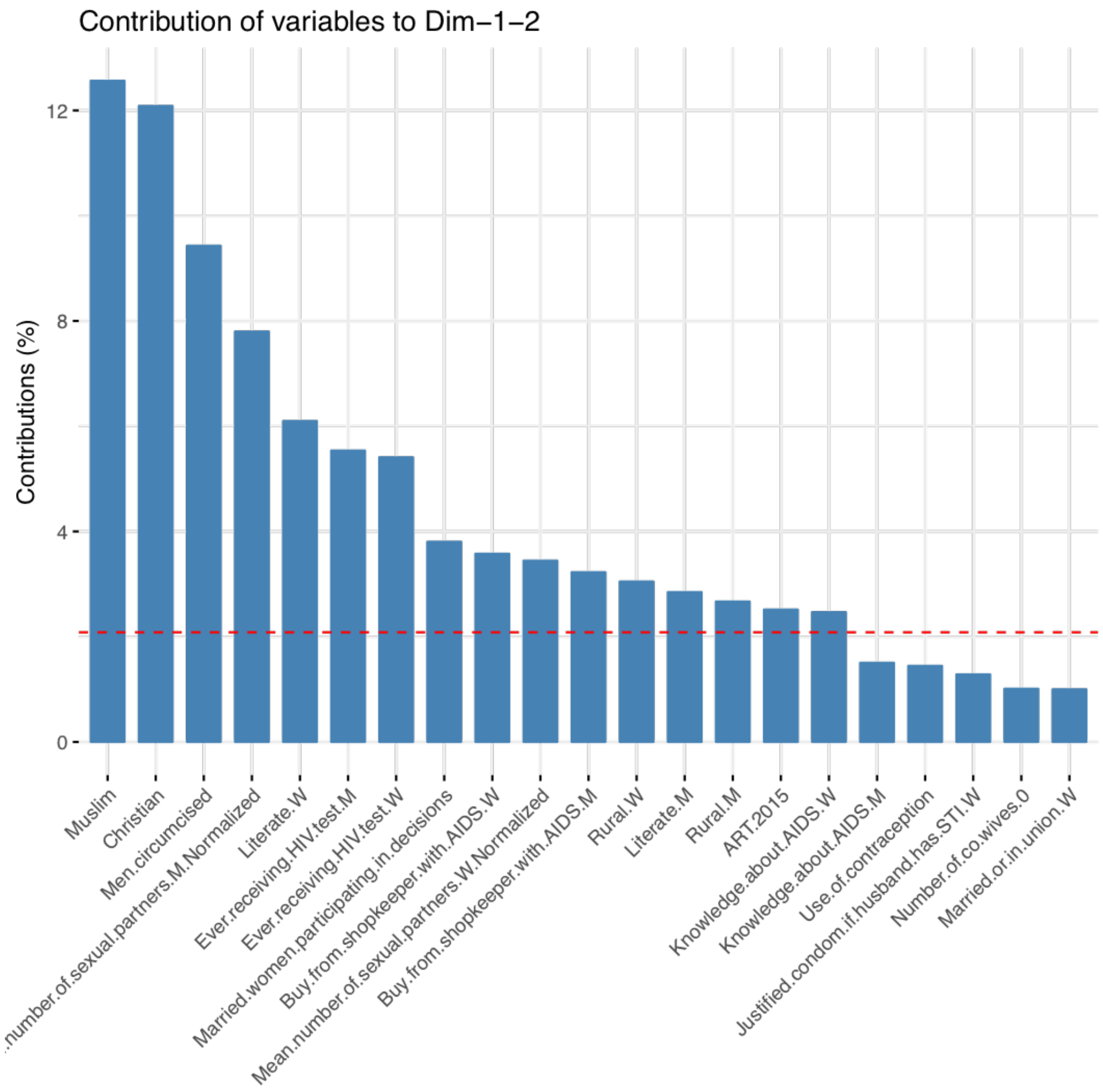


### Similarity/Dissimilarity measure

Given that each country is described by an $n$-dimensional vector, the dissimilarity $d_{i,j}$ between two countries $i$ and $j$ is defined using the Euclidian distance:

**Equation S1 - Dissimilarity measure between countries.**

$$d_{i,j} = \sqrt{\sum_{k=1}^{n} \left(c_{i,k} - c_{j,k}\right)^2},$$

where $n = 48$ is the total number of variables used in our analysis to describe a country. $c_{i,k}$ and $c_{j,k}$are the $k^{th}$ element of the n-dimensional vectors $c_i$and $c_j$, respectively.

The similarity measure $s_{i,j}$ between countries $i$ and $j$ is then defined as follows:
**Equation S2 - Euclidian similarity measure.**

$$s_{i,j} = 1 - \frac{d_{i,j}}{\max(d_{i,j})}$$

All pairwise countries similarity (or dissimilarity) data are represented by an $m \times m$ matrix $S$ (or $D$), with $m = 29$ being the number of countries. Each row and column of matrix $S$ (or $D$) represents a country, and each element $S_{ij}$ (or $D_{ij}$) is the degree of similarity (or dissimilarity), between two countries (**Figure S2**).

**Figure S2 - Similarity matrix S=($s_{i,j}$) between countries.**

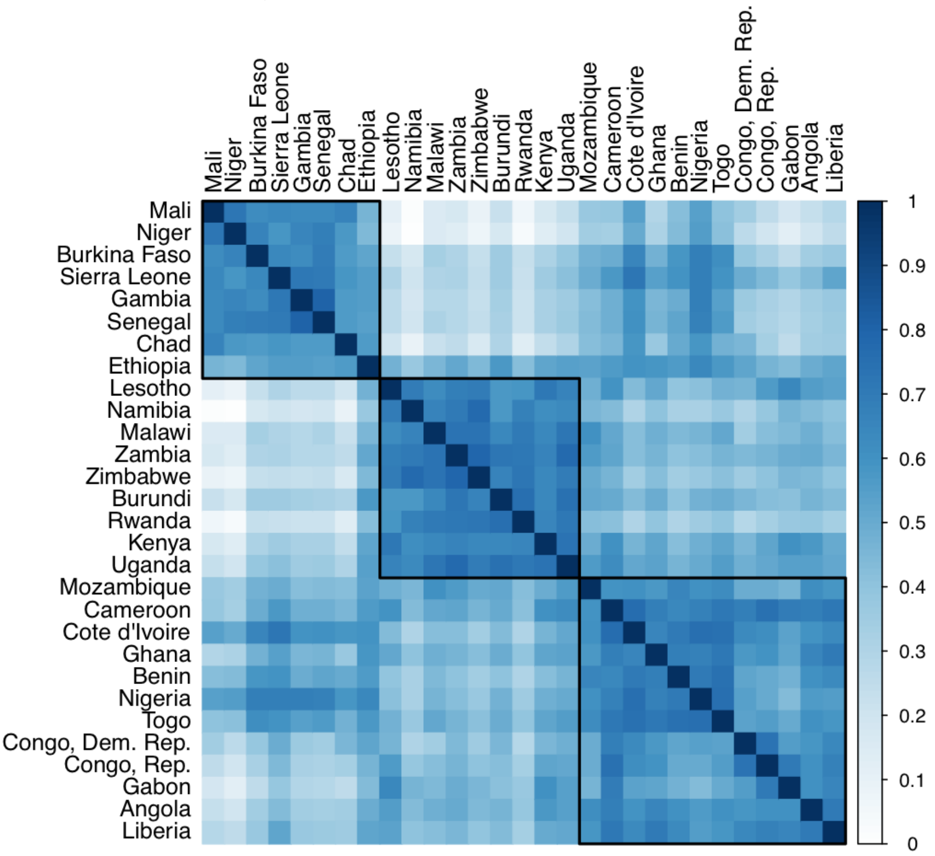


### Silhouette index

The Silhouette index is used to measure the quality of a clustering. It considers optimizing both cluster compactness and separation by minimizing the inner distance between elements of the same cluster and maximizing the distance between elements of different clusters.
For each observation (i.e. country) $c_i$, the *silhouette width* $sil(c_i)$ is defined as follows:
**Equation S3 - Silhouette width.**

$$sil(c_i) = \frac{b(c_i) - a(c_i)}{max\big(a(c_i), b(c_i)\big)},$$

where:

- $a(c_i)$ is the mean dissimilarity between $c_i$ and all other points (i.e. countries) of the cluster to which $c_i$ belongs, and
- $b(c_i) = d(c_i, C_{closest}) = \min_C d(c_i, C)$ is the dissimilarity between $c_i$ and its closest cluster $C_{closest}$, with $d(c_i, C)$ being the mean distance from $c_i$ to all observations of a cluster $C$ to which it does not belong.

The silhouette index is then obtained by averaging the silhouette widths over the whole data set:

**Equation S4 - Silhouette Index.**

$$SI = \sum_{i=1}^{m} sil(c_i),$$

where $m = 29$ is the total number of countries included in the analysis.
